# Trapped in translocation – Stalling of XPD on a crosslinked DNA substrate

**DOI:** 10.1101/2024.02.20.581127

**Authors:** Jochen Kuper, Tamsanqa Hove, Sarah Maidl, Florian Sauer, Maximilian Kempf, Elke Greiter, Hermann Neitz, Claudia Höbartner, Caroline Kisker

**Affiliations:** Rudolf Virchow Center for Integrative and Translational Bioimaging, University of Würzburg, Germany; Institute of Organic Chemistry Universität Würzburg, Am Hubland, 97074 Würzburg, Germany; Center for Nanosystems Chemistry (CNC), Universität Würzburg, 97074 Würzburg

**Author notes:** authors contributed equally (Jochen Kuper, Tamsanqa Hove).

## Abstract

The super family 2 (SF2) helicase XPD is a central component of the general transcription factor II H (TFIIH) which is essential for transcription and nucleotide excision DNA repair (NER)^1^. Within these two processes XPDs helicase function is vital for NER but not for transcription initiation, where XPD only acts as a scaffold for other factors ^2^. We deciphered one of the most enigmatic steps in XPD helicase action: the active separation of dsDNA and its stalling upon approaching an interstrand crosslink, one of the most severe DNA damages in the cell, using cryo EM. Furthermore, the structure clearly shows how dsDNA is separated and reveals a highly unusual involvement of the Arch domain in active dsDNA separation. Combined with mutagenesis and biochemical analyses, we identify distinct functional residues important for helicase activity. Surprisingly, those areas also affect core TFIIH translocase activity, revealing a yet unencountered function of XPD within the TFIIH scaffold. Importantly, our structure provides a basis for XPD damage recognition and further suggests how the NER bubble could be formed, leading to a model for the location of the XPG nuclease relative to the excised damage.

## Main

The preservation of genetic information is constantly challenged by endogenous and exogenous agents. To protect genetic information efficient countermeasures have evolved ^3^. Out of these nucleotide excision repair (NER), a template based repair mechanism, is special due to its broad substrate specificity. Substrates can range from adducts such as acetyl amino fluorene, cisplatinum DNA crosslinks to cyclobutane pyrimdine dimers (CPD), and 6,4 photoproducts (6,4 PP) ^4,5^. NER constitutes a multi-step multi-protein cascade that can be divided into 4 distinct phases. The first comprises initial lesion recognition and also demarks the two entry points of the NER cascade ^6–8^. In transcription coupled repair (TCR) RNA polymerase 2 (RNA pol 2) becomes stalled upon the encounter with a lesion. In global genome repair (GGR) the XPC complex consisting of XPC, RAD23 and Centrin 2 constantly scans the genome. Once a lesion is encountered in TCR or GGR phase two is initiated with the recruitment of TFIIH unifying the two entry branches. TFIIH is a ten subunit complex that can be divided into core TFIIH (XPB, XPD, p62, p52, p44, p34, and p8) and the cyclin activating kinase complex (CAK: consisting of CDK7, MAT1, and Cyclin H). Initially TFIIH, mainly driven by the XPB translocase, opens the bubble around the lesion. This is aided by the arrival of XPA, enhancing XPBs activity and releasing CAK from TFIIH, thereby activating XPD and initiating phase 3, the lesion verification step ^8^. Here the helicase activity of XPD further opens the bubble and when XPD encounters a lesion it becomes stalled, demarking a point of no return. Phase 4 is initiated with the 5’ phosphodiester incision by the XPF/ERCC1 nuclease complex positioned by XPA and TFIIH. Subsequently, gap filling DNA synthesis powered by PCNA, RCF and DNA polymerase δ is commenced and triggers 3’ incision of the XPG nuclease which is associated early on with the XPD subunit of TFIIH ^7^. Defects in NER can lead to severe diseases such as xeroderma pigmentosum (XP), trichothiodystrophy (TTD), and Cockayne syndrome (CS). The hallmark of XP is UV light sensitivity with a highly increased incidence towards skin cancers, whereas TTD and CS patients suffer from mental retardation premature aging and photosensitivity ^9^. Recent structural advances on higher order NER protein complexes have greatly advanced our knowledge how core TFIIH engages with undamaged Y-fork DNA and how the XPC complex and XPA in GGR or RNA pol 2, CSA, CSB, and UVSSA in TCR synergize to prepare TFIIH for bubble opening and lesion verification ^10–13^. Overall, these structures elucidated the initial recruitment of TFIIH and how XPD and XPB interact with ssDNA or dsDNA, respectively. These studies established how core TFIIH interacts with Y-forked DNA structures at the 5’ side of the incision bubble. However, vital information of how XPD and thus core TFIIH interacts with Y-forked DNA structures at the 3’ prime side of the bubble and how a damage is encountered is entirely missing so far.

We determined cryo EM structures of an XPD/p44/p62 complex from *Chaetomium thermophilum* in the presence of a Y-forked DNA structure containing an engineered interstrand crosslink at 3.1 Å resolution. Combined with functional analyses, our data show how XPD engages with DNA in the unwinding cycle revealing an unusual double active DNA opening mechanism placing the arch domain of XPD as the central player. Furthermore, we identified an unexpected role of the XPD arch domain for TFIIH translocase activity. Most importantly, our data reveal how XPD is approaching DNA damages during the unwinding cycle and how stalling occurs on interstrand crosslinked DNA, leading to a unified model for the excision bubble and how damage verification can be achieved in NER.

## XPD/p44/p62 crosslinked DNA complexes

We heterologously expressed and purified all core TFIIH subunits from *C. thermophilum* as previously described ^14^. For DNA complex formation we used equimolar amounts of XPD and the p44/p62 complex resulting in the hetero-trimeric XPD/p44/p62 complex (XPD complex) at 10 µM concentration. As DNA substrate we used a Y-fork substrate with an interstrand crosslink positioned 5 bases downstream into the dsDNA region from the unpaired junction (Fig. 1a, for details see methods section). We have shown previously that DNA containing a precursor of this crosslink represents a *bona fide* substrate for the XPD complex but in its crosslinked form can only be unwound until the crosslink is encountered ^15^ (Fig 1 a). Protein and DNA were mixed at a molar ratio of 1:1.25 and ATP was added to the mixture to initiate unwinding of the substrate. Samples were incubated for 10 min at room temperature and then vitrified for cryo EM data collection. Aqer data collection and processing (see extended Data Fig. 1) we obtained two volumes (class 1, class 2) of the XPD-DNA complex at 3.1 and 3.4 Å resolution, respectively (Fig. 1b, d and extended Data Fig. 2a-d). Both have in common that XPD and the N-terminal vWA domain of p44 could be readily built but p62 and the C-terminal zinc finger domains of p44 were unresolved in the density. This is most likely due to the high flexibility of p62 without p34, the latter serving as an additional anchor within core TFIIH ^16^ and has also been observed previously with core TFIIH in the presence of DNA ^10^. We built the model using class 1; model and data statistics are provided in Table 1. One long single stranded part of the DNA substrate extending between HD1 and HD2, is passing the iron sulfur cluster domain and prolongs through the unique XPD pore feature is clearly visible and represents the translocating strand in 5’-3’ direction (Fig. 1 b,c,d). Aqer leaving the pore this strand leads into one turn of dsDNA (Fig. 1c). The 3’-5’ end of this dsDNA extends to the ss/dsDNA junction with 3 single strand bases that separate into ssDNA at the arch domain (Fig. 1c, e). Overall, we observed 18 bases of the translocating strand and 8 bases of the non translocating strand including the crosslink (extended Data Fig. 2e). The dsDNA opening area is located at arch α1 and α4 also revealing the location of the DNA crosslink adjacent to theses helices, indicating that the XPD complex has unwound the dsDNA until it encountered the crosslink (Fig. 1e). This is further supported by the sequence assignment of the translocating strand and our observation of ADP in the ATP binding pocket suggesting that ATP hydrolysis took place (Fig. 1c and extended Data Fig. 2f). Interstrand cross linked DNA represent verified NER substrates ^17^ and the cyclobutene ^Phe^dU dimer crosslink is reasonably defined in the density. The two phenyl rings, however, could not be visualized in the sharpened density, probably due to lower resolution in that region caused by the higher flexibility of the DNA and arch domain (extended Data Fig. 2 b,e). The so called plug region of eukaryotic XPD is disordered ^10^. Class 2 mainly revealed similar features but could only be resolved to a resolution of 3.4 Å. The dsDNA region exhibits higher mobility and less features in the density. One significant difference between the two classes, however, is an extended density, that can be attributed to arch α2, which seems to elongate this helix significantly as compared to class 1 and other DNA containing core TFIIH XPD structures ^10,12^. This extension could indicate where the so called plug feature of XPD is moving in the presence of DNA (Fig. 1 b,d).

**Fig. 1:**
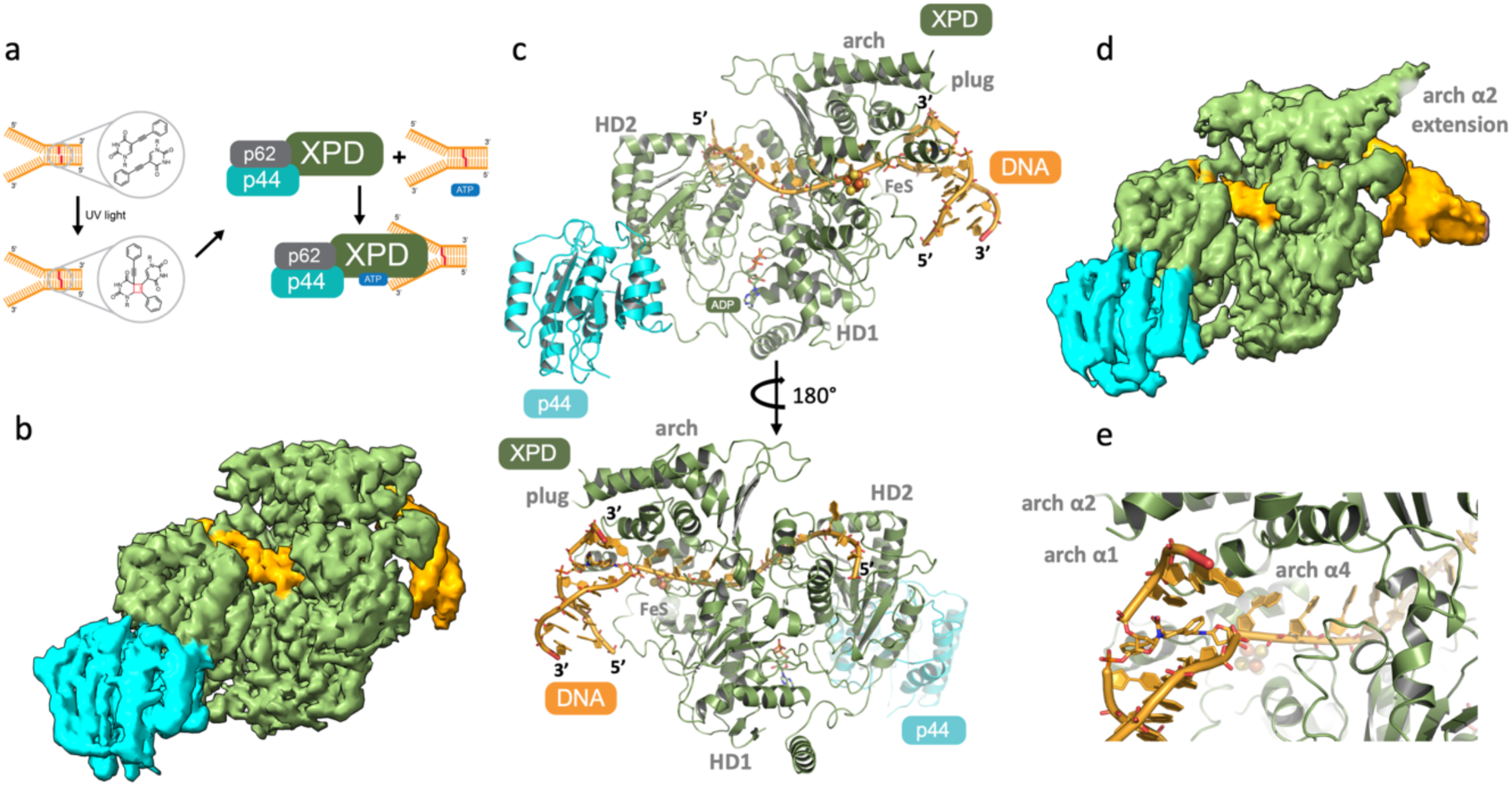
Cryo EM structure of the XPD complex in the presence of a Y-fork DNA substrate containing an engineered crosslink. **a)** Schematic description of sample preparation prior to vitrification. **b)** Cryo EM map of the class 1 XPD/p44/DNA complex. XPD is colored in green, p44 in cyan, and the DNA in orange. **c)** Structural model of the XPD/p44/DNA complex in cartoon representation. Color coding as in b. The lower panel represents the model turned 180° around its y-axis. **d)** Cryo EM map of the class 2 XPD/p44/DNA complex. Color coding as in b. **e)** Closeup on the ds/ssDNA junction and the crosslink at the arch domain of XPD. The crosslink is shown in stick representation. Color coding as in b.

**Fig. 2:**
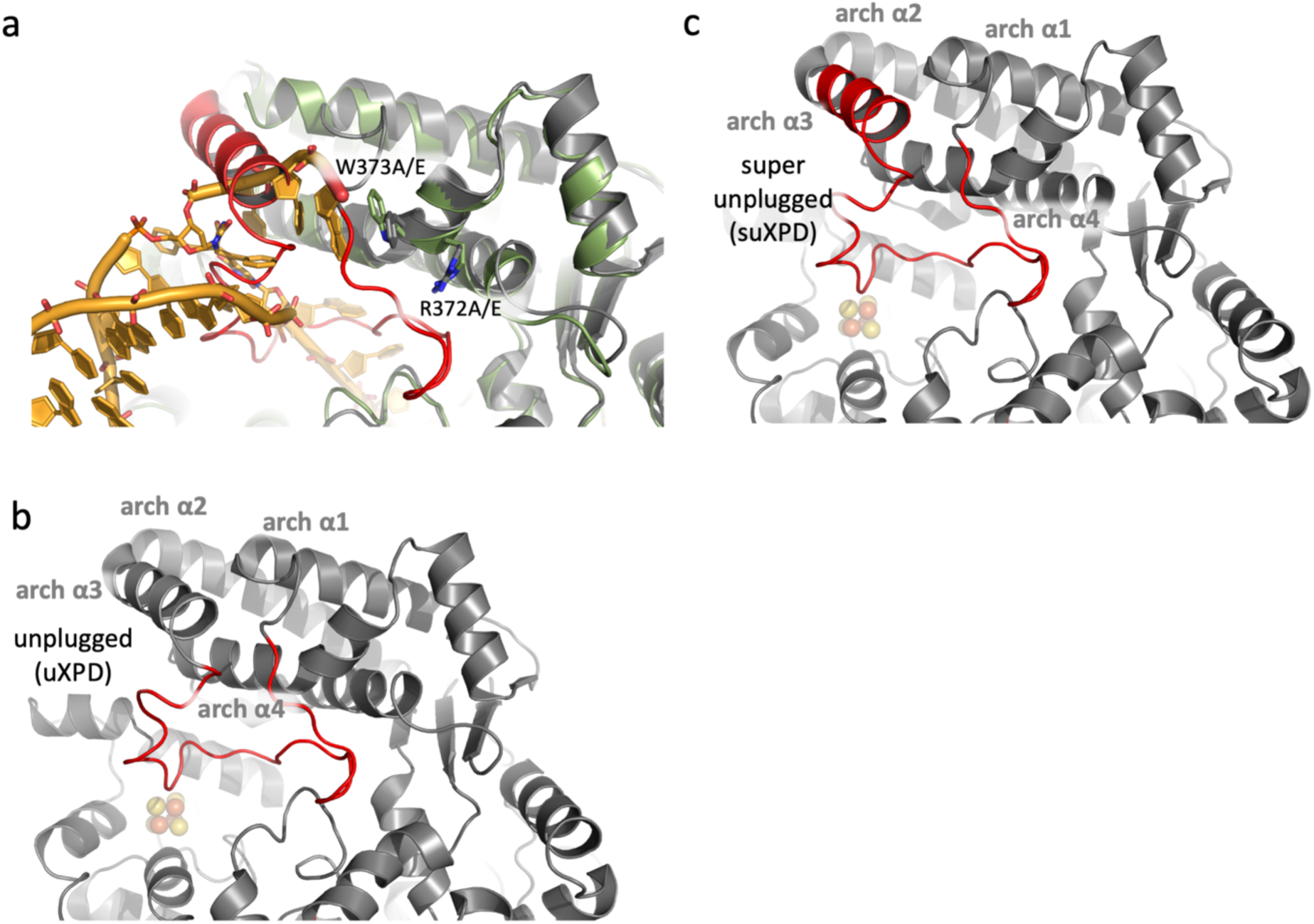
Structure based funcHonal mutagenesis. **a)** Superposition of the XPD arch domain in complex with crosslinked DNA and apo XPD (pdb entry 6nmi) in cartoon representation. Apo XPD is colored in gray with the plug region colored in red and XPD from this work is colored in green. Conserved residues subjected to mutagenesis are shown in sticks representation. **b)** Apo XPD as in a. The loop region that has been deleted to create uXPD is shown in red. 4 amino acids (STGS) to bridge the gap were inserted instead. **c)** As in b with the difference that suXPD was generated with the additional removal of α3 and the insertion of three residues (SGS) to fill the gap.

**Table 1:**
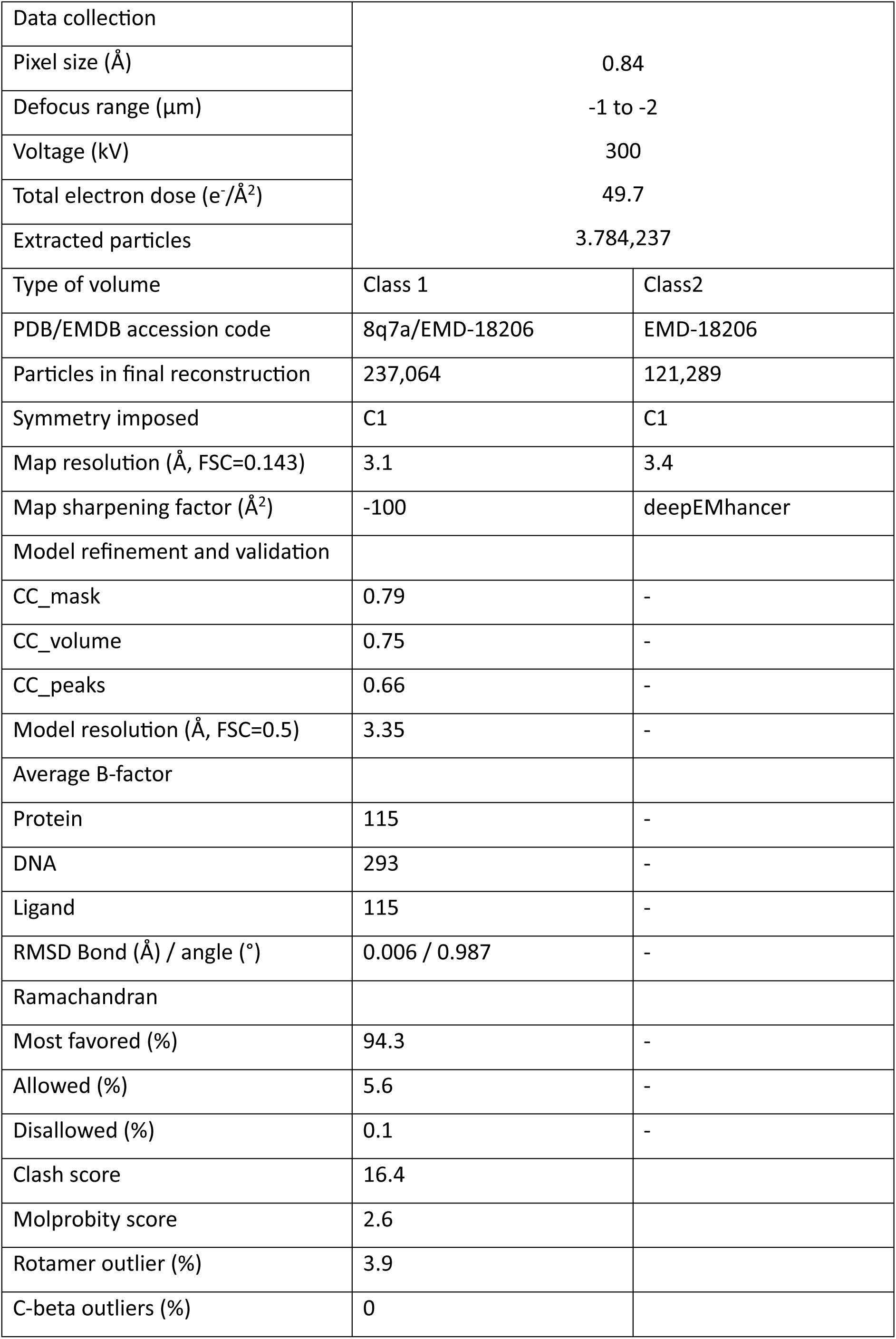
Data and model statistics.

## Functional characterization of arch domain elements

Our structure suggests that arch α4 could contribute to unwinding of the dsDNA by protein DNA interactions. We therefore mutated the conserved residues W373 and R372 located in arch α4 individually to alanine und glutamate resulting in variants W373A/E and R372A/E (Fig. 2a). W373 and R372 could be potentially involved in base and backbone interactions, respectively, although the DNA in our structure can only be observed to W373. In addition, we wanted to investigate the plug element. We therefore generated two versions of XPD in which we deleted different parts of the plug. For the so called unplugged XPD (uXPD) we removed the loop region (residues 292 to 315) and for the super-unplugged XPD (suXPD) we additionally removed most of arch α3 (residues 281 to 315) (Fig. 2b,c). All resulting variants were purified to homogeneity and thermal stability analysis confirmed correct folding of all variants comparable to wild type XPD (Fig. 3a). We subjected all variants to a detailed biochemical analysis, i.e.DNA binding, ATPase activity, and helicase activity (Fig. 3, Table 2 and extended Figure 3). A 5’ overhang hairpin substrate was used for the interaction with DNA, which binds with high affinity to wild type XPD (K_D_ 30 nM). The K_D_ values of the variants range from 27 nM to 54 nM indicating no significant influence on DNA binding (Fig. 3b). Wild type XPD displayed a robust ATPase activity of 16 µM ATP*min^-1^ in the presence of Y-fork DNA and p44. For the W373A and the R372A variants the activity is similar to wild type XPD (14 and 12 µM ATP*min^-1^, respectively) and for the W373E and R372E variants a minor decrease can be observed (each showing 10 µM ATP*min^-1^). uXPD and suXPD show a decreased ATPase activity with 6 and 8 µM ATP*min^-1^, respectively (Fig. 3c). Overall, our data indicate that DNA binding and ATPase activity are not or only slightly affected in the W373A/E and R372A/E variants and uXPD and suXPD show a reduction in ATPase activity.

**Fig. 3:**
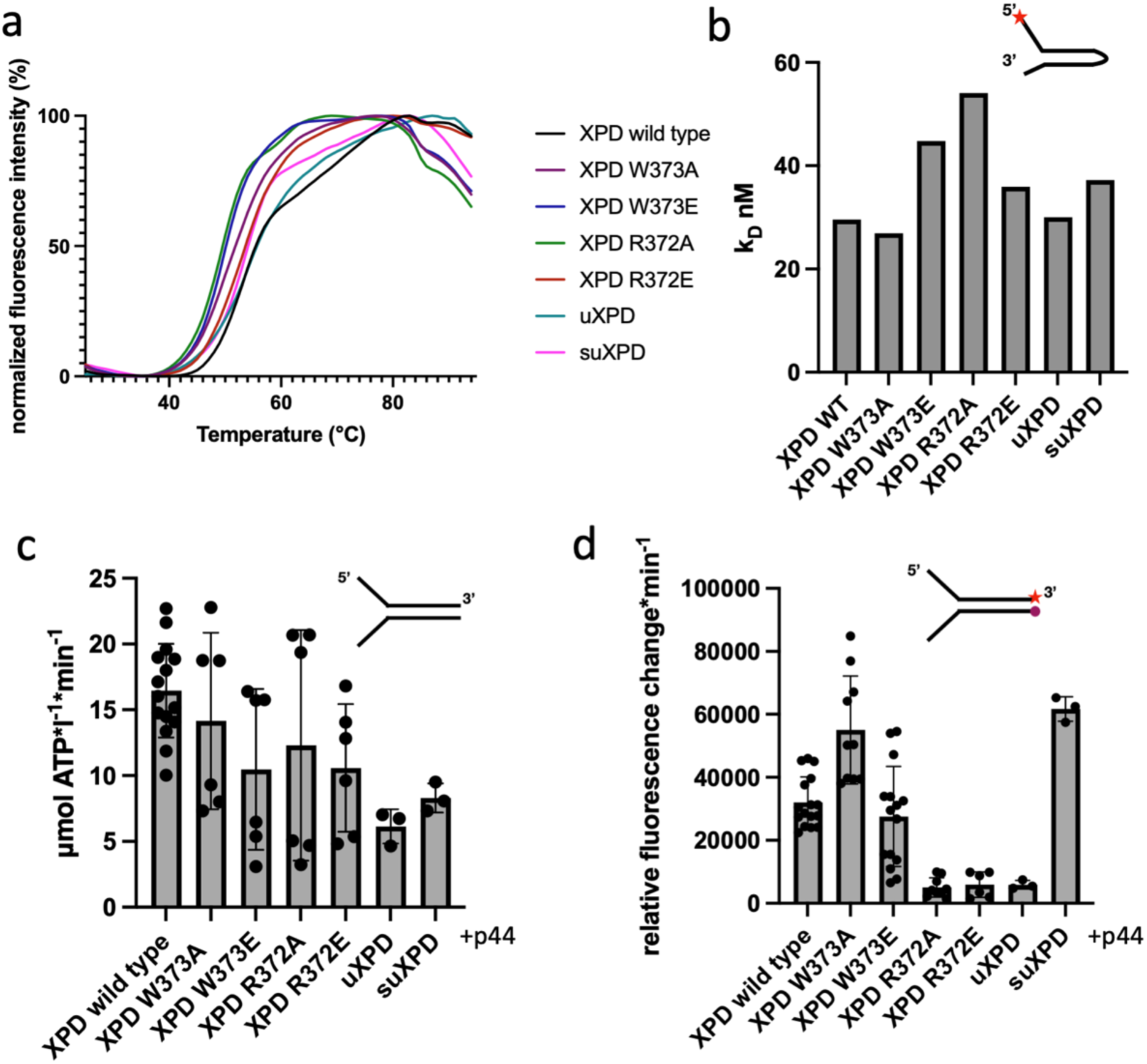
FuncHonal characterizaHon of XPD variants. **a)** Normalized thermal unfolding curves of XPD and XPD variants analyzed in this work. Melting points were derived from these curves using GraphPad Prism. **b)** Bar graph of DNA K_D_ values obtained from fluorescence anisotropy experiments using a 5’ overhang hairpin substrate. Experiments were performed in at least three technical replicates and one biological replicate (no biological replicate for uXPO and suXPD). The red star marks the Cy3 label. **c)** ATPase activity of XPD and its variants in the presence of a Y-fork substrate and p44. Experiments were performed in at least three technical replicates and one biological replicate (no biological replicate for uXPO and suXPD). **d)** Helicase activity of XPD and its variants in the presence of a fluorescence labeled Y-fork substrate and p44. The red star denotes a Cy3 label at the 3’ end that is quenched by a dabcyl moiety at the 5’ end of the complementary strand. Experiments were performed in at least three technical replicates and one biological replicate (no biological replicate for uXPO and suXPD). The data were analyzed using GraphPad Prism. All values are also listed in Table 2.

**Table 2:** Biochemical data.

| Protein | DNA<br>binding kD<br>(R<sup>2</sup> of fit) | ATPase<br>$\mu\text{M} \cdot \text{min}^{-1} \pm$<br>SD | Helicase<br>$\Delta F \cdot \text{min}^{-1}$<br>$\cdot 1000 \pm \text{SD}$ | Melting point<br>°C | Translocase<br>$\Delta F \cdot \text{min}^{-1} \cdot 1000$ |
| --- | --- | --- | --- | --- | --- |
| XDP wild type | 30 (0.97) | 16 ± 4 | 32 ± 8 | 56 | 2.1 ± 0.9 |
| R372A | 54 (0.84) | 12 ± 8 | 5 ± 3 | 49 | n.d. |
| R372E | 36 (0.88) | 10 ± 5 | 6 ± 4 | 54 | n.d. |
| W373A | 27 (0.91) | 14 ± 7 | 55 ± 17 | 51 | n.d. |
| W373E | 45 (0.84) | 10 ± 6 | 28 ± 16 | 50 | n.d. |
| uXPD | 30 (0.98) | 6 ± 1 | 6 ± 1 | 56 | 1.3 ± 0.6 |
| suXPD | 37 (0.99) | 8 ± 1 | 62 ± 4 | 54 | 6.0 ± 2.7 |

Significant differences between the wild-type protein and the variants, however, were observed in our helicase assay utilizing a Y-fork substrate. W373A showed enhanced activity that amounts to 172 % of wild type activity, whereas the glutamate variant, W372E, displayed only a minor decrease to 88 % of wild type activity. Both R372 variants showed a strong decrease in helicase activity (16 and 19 %) indicating they are highly relevant for dsDNA separation. uXPD also displayed a significant decrease in helicase activity (19%) whereas the additional removal of arch α3 led to the highest helicase activity with 194 % of the wild type level (Fig. 3d) despite its lower ATPase activity.

## The unusual role of the arch domain for XPD helicase action

Our data clearly indicate that the arch domain is essential for XPD helicase action (Fig. 4). The total removal of the plug region (suXPD) led to a hyperactive helicase, whereas removing only the loop region impairs activity (uXPD). In our DNA bound structure, the plug region is disordered and thus we could not observe any interaction of the plug with the double stranded part of the DNA. Our class 2 data, however, clearly show an elongation of arch α2 that could be attributed to arch α3 moving up and “fusing” with α2 when DNA is bound (Fig. 4b). To accommodate this move, the entire loop region of the plug has to undergo a conformational change to move with the helix upwards. Removal of the loop disables this movement and arch α3 stays in position thereby hindering helicase activity and thus strengthening the notion of the plug acting as a negative regulatory element ^10^. Interestingly, arch α3 and the plug are shielding arch α4 in which we previously located functionally important residues for helicase action. Activating XPD therefore requires the plug to move upward and out and could possibly mediate other interactions for instance with XPG ^10^ (Fig. 4b).

**Fig. 4:**
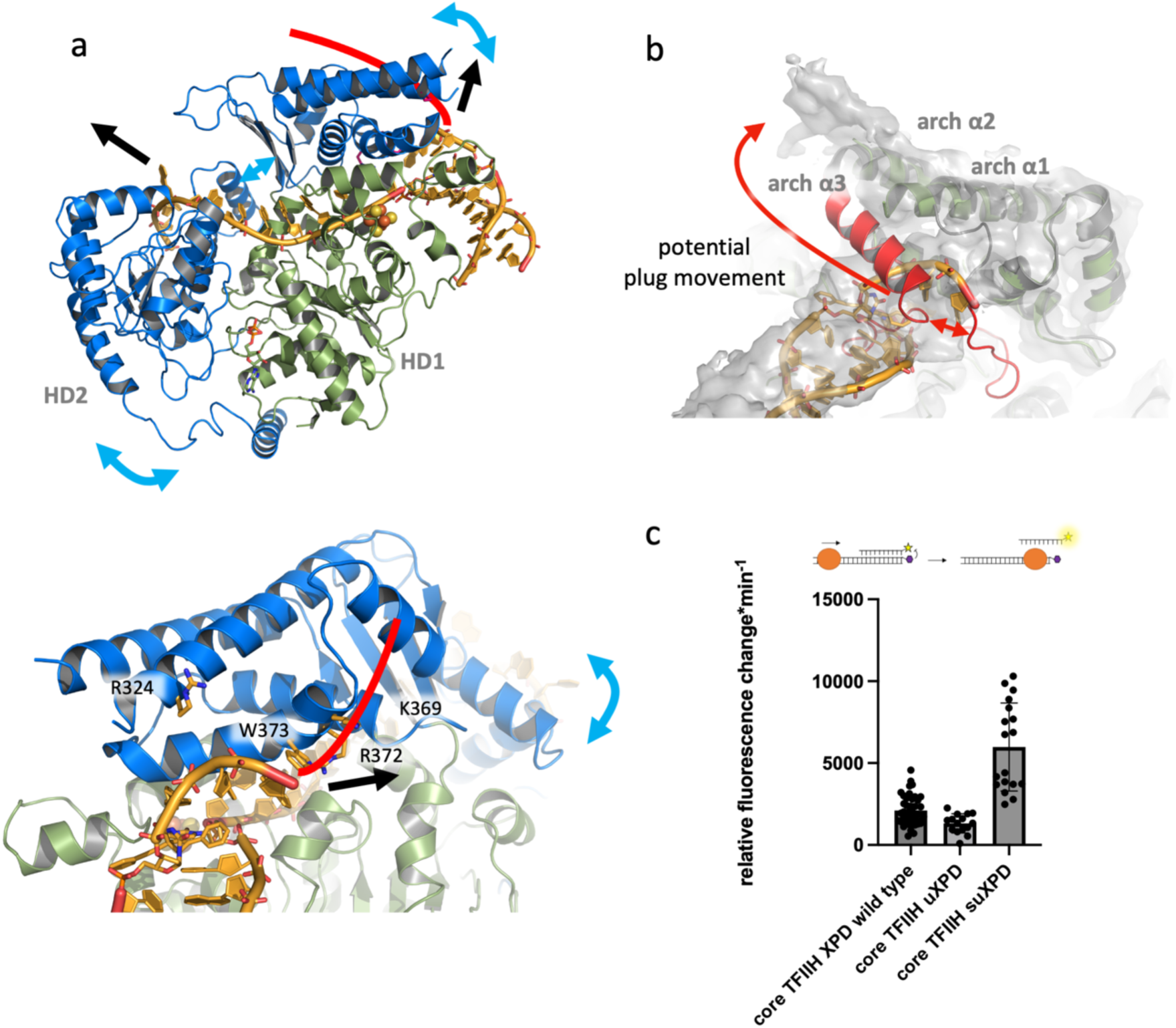
XPD helicase mechanism and plug dynamics. **a)** Two views of the structure of XPD in complex with the crosslinked DNA substrate in cartoon representation. HD2 and the arch domain are colored in blue and the remainder of XPD is colored in green. DNA is shown in orange. Blue arrows indicate the possible domain movement during ATP hydrolysis and ssDNA translocation. Black arrows indicate the direction of the DNA movement. The red line indicates the likely path of the non-translocated strand along the arch domain supported by functional data. **b)** Possible movement of the plug based class 2 density that suggests the elongation of arch α2 by the plug region. The long red arrow indicates the motion of the plug whereas the short red double arrow shows why the movement could be hindered in uXPD. **c)** Translocase activity of XPD and plug variants determined using a triplex disruption assay. Experiments were performed in at least three technical replicates and one biological replicate. The data were analyzed using GraphPad Prism. All values are also listed in Table 2.

With the plug in an out position XPD is now primed for helicase activity. It has been established for the bacterial XPD homolog DinG that ATP dependent ssDNA translocation mediated by HD1 and HD2 movement is translated into a swingout motion of the arch domain via a pusher helix in HD2 ^18^(Fig. 4a). Based on our structure the proposed swingout of the arch domain would lead to a highly unusual active pulling on the non translocated strand that is separated at the arch domain DNA junction (Fig.4a). This mechanism would actively engage both the translocating and non-translocating strand to promote the separation of dsDNA. This hypothesis is further supported by the presence of essential residues that modulate helicase activity (Fig. 4a, lower panel and ^19^). Our analysis shows that R372 is vital for activity, possibly due to phosphate backbone interactions whereas W373 seems to be involved in base interactions, which, if removed, increases DNA separation capacity. In a previous study, we identified R324 and K369 to be essential for helicase activity ^19^. R324 is located in close vicinity to the phosphate backbone of the non-translocating ssDNA strand and K369 lies in the proposed path for the non-translocating strand across the arch domain (Fig. 4). Importantly, this proposed mechanism is most likely conserved in all XPD homologs including human RTEL, FANCJ, and DDX11 since they all contain arch domains that could interact with DNA in a similar way (extended Data Fig. 4)

## A novel role for XPD in the TFIIH translocase complex

A superposition of our XPD/p44 DNA structure with core TFIIH reveals no significant differences in the ssDNA interactions with respect to the translocated strand (Fig. 5a). Furthermore, the overall orientation of XPD/p44 is highly comparable to that in core TFIIH (1.7 Å rmsd) showing that our structural and functional interpretations can be readily transferred to core TFIIH.

**Fig. 5:**
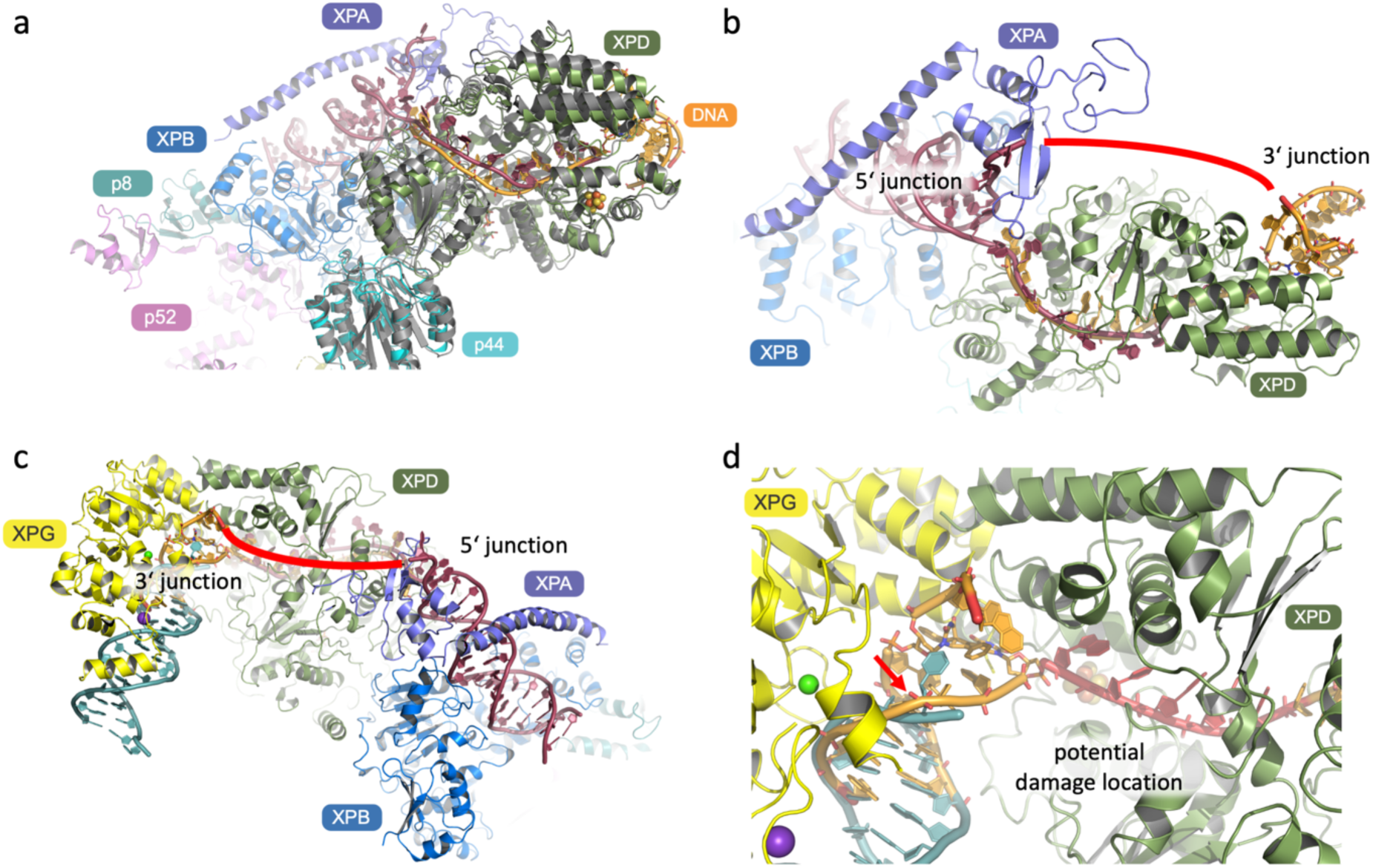
Model of the early NER incision bubble. **a)** Superposition of the XPD/p44/DNA complex (color code as in Fig. 1c) from this work with XPD/p44 (colored in grey) in core TFIIH bound to DNA (pdb entry 6ro4). The DNA from 6ro4 is colored in dark red. **b)** Model of the early incision bubble of core TFIIH. The XPD structure is from this work. Both DNAs are visible and could form the early bubble. The unresolved part of the DNA is indicated by the red line. Color coding as in a). **c)** Model of XPG at the 3’ junction based on the superposition of the XPG substrate DNA (cyan) with the dsDNA of the XPD/p44/DNA complex. **d)** Closeup of c) indicating a possible incision site (red arrow). Note that the crosslink forces the DNA to be closed at the potential incision site. Non crosslink damaged DNA could be already separated at that position. Bases that could represent the position of a damage based on the average excised fragment are colored in red.

Prior to our plug analysis, all data indicate that XPD is not involved in the translocase activity of core TFIIH ^14,20^. To our surprise, however, we observed that uXPD and suXPD modulate core TFIIH translocase activity. uXPD displays a moderately decreased activity of 62 % compared to wt XPD whereas the suXPD increased the activity of core TFIIH to 286 % of wild type indicating a strong influence of this region on core TFIIH translocase (Fig. 4c). This observed boost upon plug removal likely has implications for NER but not for the transcriptional TFIIH translocase where holo TFIIH is involved and XPD is inhibited by the CAK complex. ^2,20–22^. In contrast, additional translocase activity of core TFIIH might be required in repair functions. It has been proposed that TFIIH could push back RNA pol 2 prior to damage verification in TCR ^23,24^. During this action a more active translocase might be necessary to facilitate the push back and engage core TFIIH with damage location. This is supported by the observation that repair factors like XPA and XPG also increase core TFIIH translocase and helicase activity ^10,25^. XPA could facilitate CAK removal, thus enhancing plug flexibility. The flexible plug could more readily permit the arch domain interaction with XPG ^19^ resulting in an open plug conformation as indicated in the class 2 data (Fig. 4b) which is essentially mimicked by suXPD that also shows enhanced translocase and helicase properties.

## Implications for damage recognition and NER bubble formation

Our data provide insights on the formation of the NER bubble that needs to be formed prior to double incision (Fig. 5b). A merge of our structure with the core TFIIH-XPA-DNA structure (^10^, pdb code: 6ro4) leads to a stretch of 11 bases ssDNA spanning the XPD helicase which is framed on both sides by the two dsDNA junctions. The proposed route of the non-translocating strand bridges the two ends of both structures where the DNA could not be resolved due to high flexibility. This model likely represents an early stage of bubble opening by the XPB/XPA translocase activity ^12^ directly followed by XPD engagement and initial unwinding. This initial bubble would then be widened prior to the primary 5’ incision since both endonucleases cut at a ssDNA/dsDNA junction ^6^. The excised fragment spans on average 27 bases and with the damage being 5-6 bases away from the 3’ end ^26^, further bubble opening is required. Depending on the location of the damage, two scenarios could be envisioned. In scenario 1, the damage is located further away in the 3’ direction of XPD translocation so that the bubble is widened by XPD helicase activity and subsequently ssDNA is protected by RPA. XPD then reaches the damage and is stalled which delivers the signal for 5’ incision. In scenario 2, the damage is located close to the 3’ area of XPD translocation and XPD is stalled on the damage early. In this case XPB/XPA activity is required to extend the bubble with the support of RPA. Bubble size restriction in this case might be induced through distance limitations in the unwinding process due to the core TFIIH structural framework. In either case, NER would proceed with 5’ incision followed by 3’ incision.

The location of the dsDNA part of the substrate enables us to model a possible engagement of XPG to perform the 3’ cut (Fig. 5c). We superimposed the dsDNA part of our structure with the DNA bound structure of the yeast XPG homolog Rad2 ^27^ which leads to a complex with the catalytic core of Rad2 being closely located at the DNA junction thus revealing a potential 3’ cuyng site (Fig. 5d). This site is not yet at the ss/dsDNA junction in our structure. However, due to the presence of the chemical crosslink, further opening of the DNA is inhibited and might occur more likely when non interstrand crosslink damages are encountered. Based on this position, possible damage locations in the 5’ direction are marked which all fall into regions that have been previously implicated in XPD damage verification ^18,28,29^ strongly supporting our model.

Our data provide vital information on the essential XPD helicase but also the other iron sulfur cluster containing helicases FANCJ, RTEL1 and DDX11 revealing an unusual double active unwinding mechanism engaging both strands. Furthermore, we identified the XPD plug as a regulatory element that is not only involved in helicase function but also actively modulates core TFIIH translocase activity with implications for TCR. Lastly, we provide a model of the early NER incision bubble that suggests a mechanism for XPG engagement in incision.

## Extended Data Figures

**Extended Data Fig. 1:**
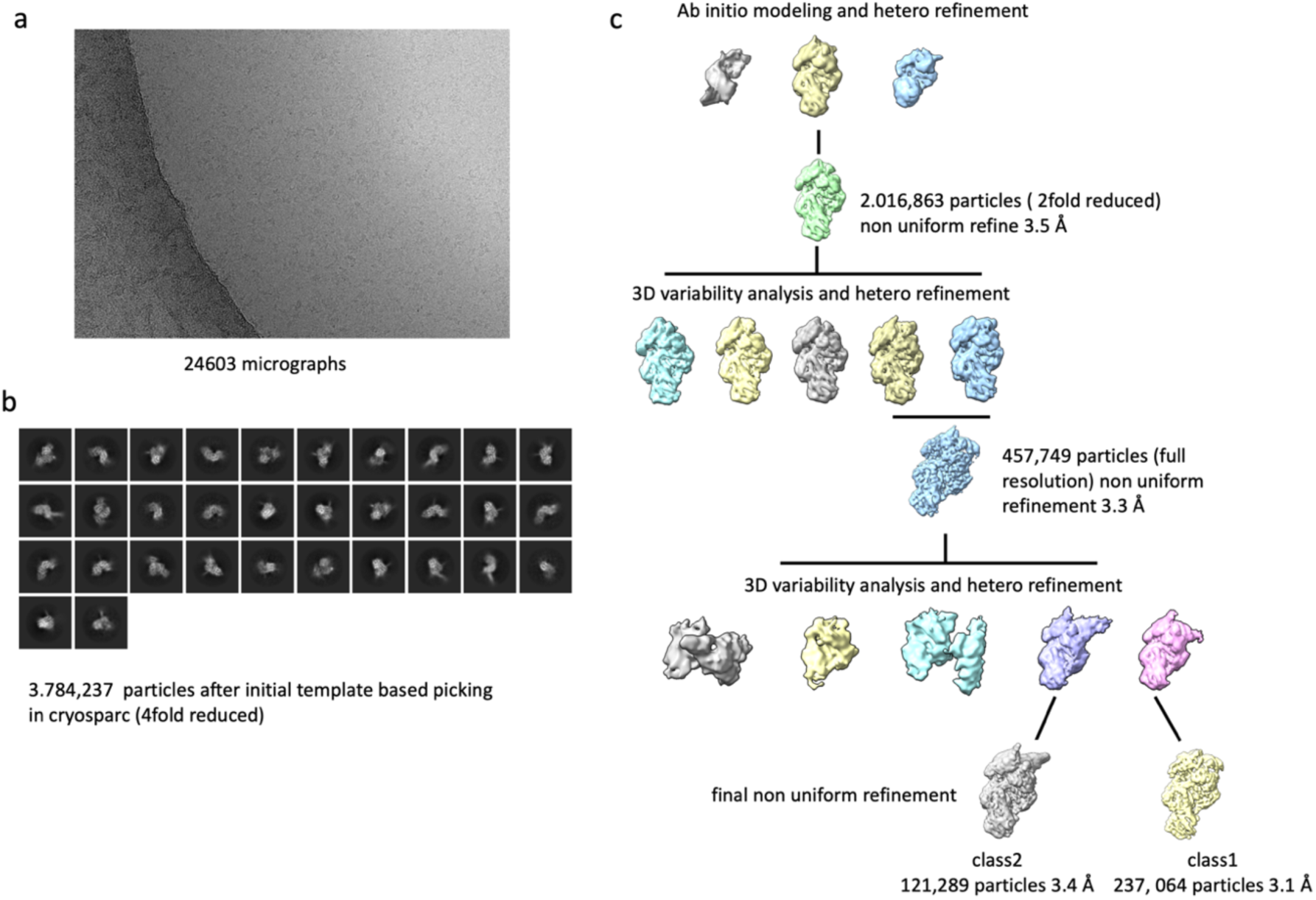
Workflow of cryo EM data processing. **a)** Representative micrograph of the XPD complex/DNA sample revealing the single particles to be evenly distributed. **b)** Reference free 2D class averages obtained with cryosparc. The selected classes have been obtained from the initial round of template based particle picking and represent 3.784,237 particles. **c)** Schematic workflow of data processing in cryosparc. All employed steps are indicated.

**Extended Data Fig. 2:**
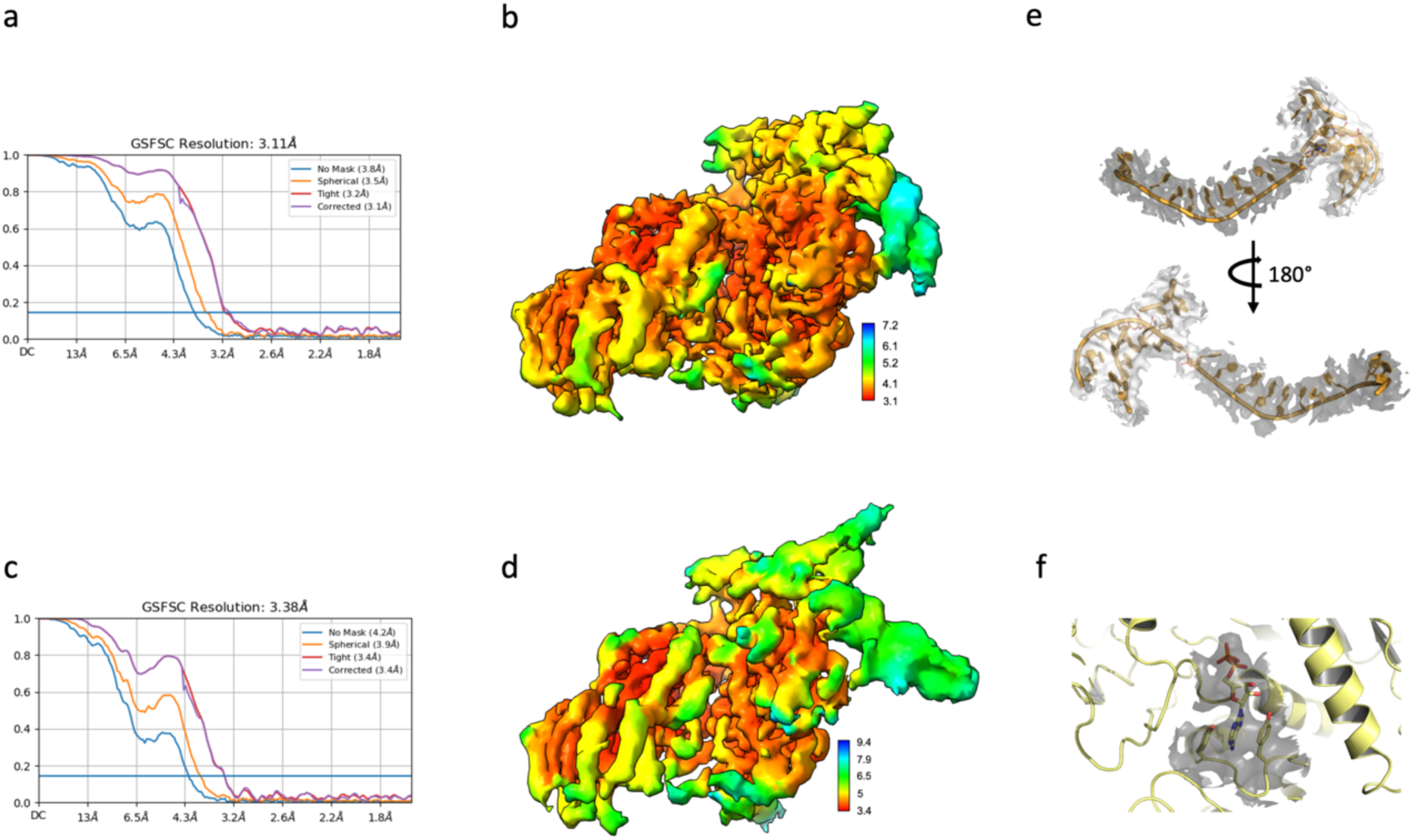
Map quality of the resulting cryo EM maps. **a,c)** Gold standard Fourier shell correlation plots for class 1 (a) and class 2 (c) indicating the resolution limits of the data. **b,d)** Local resolution maps of class 1 (b) and class 2 (d) cryo EM maps. **e)** Density of the DNA substrate. Gray density represents the map plotted with 12 sigma, light gray density represents lower resolution data of the same map plotted at 5 sigma. The DNA is shown in cartoon mode. **f)** Density for ADP plotted at 12 sigma. ADP is shown in stick mode, whereas the protein is drawn as cartoon.

**Extended Data Fig. 3:**
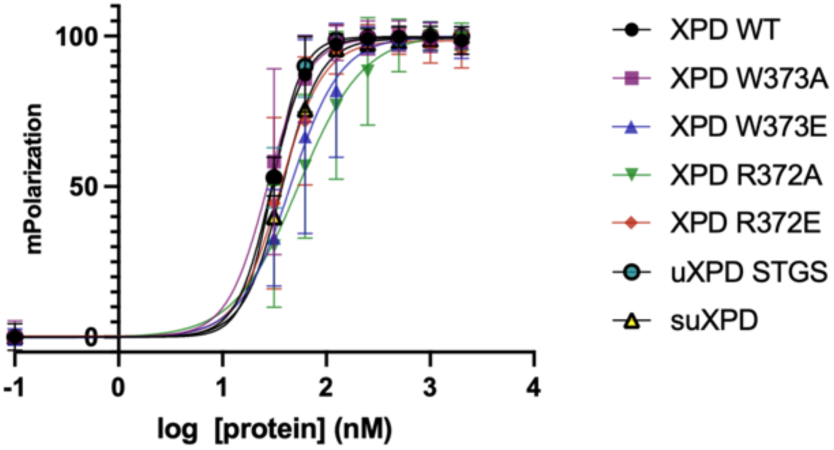
DNA binding analysis. Normalized plot of fluorescence anisotropy data used to determine the K_D_ values shown in Fig. 3b. Curves were fitted with GraphPad Prism and represent the averages of at least three technical replicates and two biological replicates (no biological replicate for uXPD and suXPD). Mean values are plotted with their associated SD.

**Extended Data Fig. 4:**
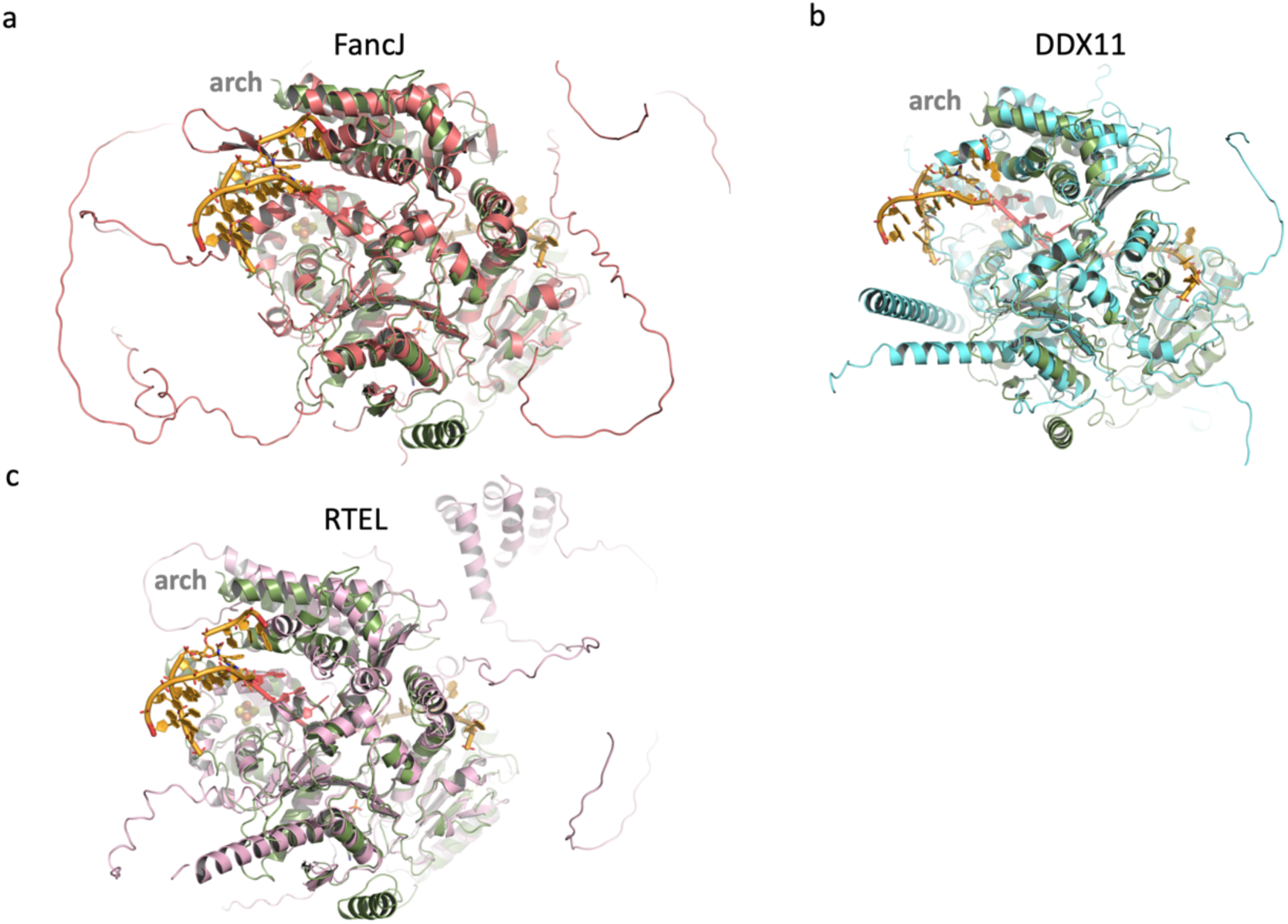
Iron sulfur cluster containing helicases. Superposition of the structure obtained in this work with AlphaFold models of **a)** FancJ (colored salmon), **b)** DDX11 (colored cyan), and **c)** RTEL (colored in pink). All models are shown as cartoon. All proteins contain an arch domain suggesting a conserved mode for unwinding DNA.

## Methods

### Protein expression, purification, and mutagenesis

All proteins were purified as described in ^14^ with the exceptions mentioned below. For the generation of the *C. thermophilum* XPD/p44/p62 complex we co-expressed and purified the p44/p62 complex and then mixed it with separately purified XPD. The N-terminal domain of p44 was produced as described in ^2^. XPD single amino acid variants were generated via site directed mutagenesis ^30^. uXPD and suXPD were generated by deleting the sequences coding for residues 292 to 315 for uXPD (replaced with the linker sequence STGS) and 281 to 315 for suXPD (replaced with the linker sequence SGS) using sequence and ligation independent cloning ^31^. All variants were purified following the protocol employed for wild type XPD without modifications.

### Cryo EM Sample preparation and data collection

5-phenylethynyl-2ʹ-deoxyuridine (_Phe_dU) containing DNA to generate the cyclobutene _Phe_dU dimer crosslink by alkene-alkyne [2 + 2] photocycloaddition was produced and annealed as previously described (Fork 1: 5’-AGCTACCATGCCTGCACGAATTAAGCA(_Phe_dU)CGCGTAATCATGGTCATAG-3’, Fork 2: 5’-CTATGACCATGATTACGC(_Phe_dU)CTGCTTGGAATCCTGACGAACTGTAGA-3‘)_15_. We mixed the crosslink containing DNA substrate with 10 µM XPD/p44/p62 complex at a molar ratio of 1.25:1. Samples were mixed in 20 mM Hepes pH 7.5, 50 mM KCl, 1 mM TCEP, and 5mM MgCl_2_, the same buffer which was also used for XPD activity analysis to ensure efficient partial substrate unwinding (see also _15_). Samples were incubated on ice for 10 min and 5 mM ATP were added to initiate the helicase cycle. The mixture was incubated further for 10 min at room temperature to allow efficient substrate engagement. Samples were subsequently immediately used for cryo-grid preparation. Three microliters of sample were applied to glow-discharged R2/2 carbon grids (Quantifoil), which were blotted for 5 s and at a blot force of 25 and plunge-frozen in liquid ethane with a Vitrobot Mark IV (ThermoFisher) operated at 4 °C and 100% humidity. Data were collected at the CM01 facility of the ESRF _32_. Micrographs were acquired at a nominal magnification of ×105,000 (0.84 Å/pixel) using a dose rate of 17.5 e^−^/pixel per s over the Mme of 2 s resulting in a total dose of 49.7 e^−^/Å^2^ fractionated over 50 frames. Movies were recorded over a defocus spread of −1 µM to −2 µM with 0.2 µM step size. Overall, a total number 24603 movies were collected.

### Cryo EM processing and model building

Motion correction and dose weighting was performed using MotionCor2 ^33^ within the cryosparc framework ^34^. CTF correction was achieved using patch CTF from cryosparc ^34^. We used the template based picking algorithm in cryosparc with a low resolution model of the complex that has been obtained in house using a ThermoFisher Titan-Krios G3 with an X-FEG source, 300 kV and a Falcon III camera. Initial picking and 2D classification (4-fold binning) in cryosparc resulted in 3.784,237 particles that were subjected to cryo sparc *ab ini3o* modeling with subsequent hetero refinement of the resulting 3 classes which revealed one class containing XPD/p44 with 2.016,863 particles that were subsequently re-extracted with a 2 fold binning. This set was subjected to non-uniform 3D refinement in cryosparc and subsequently analyzed using the 3D variability function. 3D variability analysis resulted in 5 clusters that were subjected to further hetero refinement. Of the 5 classes 2 contained dsDNA located at the arch domain that were pooled and re-extracted at full size (box size 384 pixel). Non uniform 3D refinement was performed on this set followed by an additional round of 3D variability analysis and hetero refinement. In this round the final class 1 and class 2 data were obtained with 237,064 and 121,289 particles, respectively (see extended Data Fig. 1 for details and overview). This resulted in an overall resolution of 3.1 Å for class 1 and 3.4 Å for class 2 as defined by the gold standard Fourier shell 0.143 correlation criterion (extended Data Fig. 2a,c). Local resolution maps show the highest resolution for the XPD and ssDNA parts of the density degrading in the arch domain and dsDNA regions (extended Data Fig. 2 b,d). For model building we used AlphaFold2 models of p44 and XPD from *C. thermophilum* that were assembled based on the arrangement of XPD/p44 in pdb entry 6ro4. This complex was used for map docking in Phenix ^35^. The results of the initial map docking were further improved by manual model building in coot. No further density was observed that could be attributed to p62 or the C-terminal part of p44. The DNA and the crosslink were built manually in coot and sequence assignment was based on crosslink location and map quality. Manual model building was iterated with rounds of real space refinement using the refmac5 ^36^ pipeline in ccpem ^37^ and phenix realspace refine ^35^. The final model and density correlation statistics are given in Table 1.

### *In vitro* DNA dependent ATPase activity assay

XPD ATPase activity was measured utilizing an *in vitro* ATPase assay in which ATP consumption is coupled to the oxidation of NADH via pyruvate kinase and lactate dehydrogenase activities as described previously ^19^. The assay was carried out under saturating concentrations of ATP (5 mM) using XPD wild type, variants and p44 at a concentration of 250 nM. Y-fork DNA (strand 1: 5ʹ-AGCTACCATGCCTGCACGAATTAAGCAATTCGTAATCATGGTCATAGC-3ʹ, strand 2: 5’-GCTATGACCATGATTACGAATTGCTTGGAATCCTGACGAACTGTAG-3‘) was added at final concentrations of 250 nM. The mix of all reagents, with the exception of ATP, was preincubated at 30 °C until a stable base line was achieved. Enzyme catalysis was initiated by the addition of ATP. The activity profiles were measured at 340 nm using a Clariostar (BMG labtech) plate reader. Reactions were followed until total NADH consumption was reached. Initial velocities were recorded and ATP consumption was determined using the molar extinction coefficient of NADH. Measurements were carried out with at least three technical replicates and mean values were plotted with their associated SD and two biological replicates (no biological replicate for uXPD and suXPD). Mean and SD were determined using the GraphPad Prism soqware.

### *In vitro* helicase assay

Helicase activity was analyzed utilizing a fluorescence-based helicase assay described in ^19^. We used a Y-fork substrate with a cy3 label at the 3ʹ end of the translocated strand (5ʹ-AGCTACCATGCCTGCACGAATTAAGCAATTCGTAATCATGGTCATAGC-3ʹ-cy3, and a dabcyl modification on the 5ʹ end of the opposite strand (Dabcyl-5ʹ-GCTATGACCATGATTACGAATTGCTTGGAATCCTGACGAACTGTAG-3ʹ). Assays were carried out in 20 mM HEPES pH 7.5, 50 mM KCl, 5 mM MgCl_2_, and 1 mM TCEP. DNA and protein were used at a concentrations of 250 nM. Helicase activity of wild-type XPD and its variants was measured at equimolar concentrations of XPD/p44. The mix of all reagents, with the exception of ATP, was preincubated at 30 °C until a stable base line was achieved. The reaction was subsequently started with the addition of 5 mM ATP. Live Kinetic measurements were recorded with a Clariostar plate reader (BMG labtech). Initial velocities of the kinetic data curves were fitted with the MARS soqware package (BMG labtech) and represent the averages of at least three technical replicates two biological replicates (no biological replicate for uXPD and suXPD). Mean values are plotted with their associated SD. Mean and SD were determined using the GraphPad Prism soqware.

### Fluorescence anisotropy

DNA binding was analyzed by fluorescence anisotropy employing a self annealing hairpin with a 5‘ cy3 label (Cy3-5ʹ-TTTTTTTTTTTTTTTCCCGGCCATGCGAAGCATGGCCGTT-3ʹ). Assays were carried out in 20 mM HEPES pH 7.5, 50 mM KCl, 5 mM MgCl_2_, 1 mM TCEP, and 5 nM DNA at room temperature. Wild-type XPD and variants were used at concentrations of 31.5– 2000 nM. Aqer mixing, the reaction was incubated for 5 min prior to recording. Fluorescence was detected at an excitation wavelength of 540 nm and an emission wavelength of 590 nm with a Clariostar plate reader (BMG labtech). The gain was adjusted to a well containing buffer and DNA but no protein. Curves were fitted with GraphPad Prism and represent the averages of at least three technical replicates and two biological replicates (no biological replicate for uXPD and suXPD). Mean values are plotted with their associated SD (extended Data Fig. 3).

### In vitro translocase activity assay

Core TFIIH dsDNA-translocase activity was detected using a well established triplex disruption assay ^10,20^. DsDNA-translocase activity was measured by displacement of a fluorescently labeled triplex forming oligonucleotide (TFO) from a triple helix DNA substrate and was carried out as published before ^14^ using 150 nM of triplex DNA (fw: 5ʹ-GTCTTCTTTTAAACACTATCTTCCTGCTCATTTCTTTCTTCTTTCTTTTCTT-3ʹ; rv: 5ʹ-BHQ-AAGAAAAGAAAGAAGAAAGAAATGAGCAGGAAGATAGTGTTTAAAAGAAGAC-3ʹ and a 5ʹ Cy3-tagged TFO 5ʹ-Cy3-TTCTTTTCTTTCTTCTTTCTTT-3ʹ). A black hole quencher (BHQ) was used for Cy3 quenching. Correct triplex formation was confirmed via native PAGE. The baseline was recorded for 10 to 15 minutes prior to addition of 2 mM ATP. TFO displacement was measured for 60 min at an excitation wavelength of 520–540 nm and an emission wavelength of 590– 620 nm with a gain of 1900 using a Clariostar plate reader (BMG labtech) in 384-well F-bottom FLUOTRAC™ high binding microplates (Greiner Bio-One). Core TFIIH was assembled with all subunits present in equimolar amounts with a final concentration of 500 nM. Initial velocities of the kinetic data curves were fitted with the MARS soqware package (BMG labtech) and represent the averages of at least three technical replicates and two biological replicates. Mean values are plotted with their associated SD. Mean and SD were determined using the GraphPad Prism soqware.

### Thermal unfolding assays

Correct folding of the XPD variants was tested by thermal shiq assays using sypro orange (Invitrogen) and a qPCR machine (Stratagene mx3005p). The final reaction mix of 25 μl comprised 2.5 μM XPD, 0.1% sypro orange, 20 mM HEPES pH 7.5, 200 mM NaCl, 5 mM MgCl_2_, and 1 mM TCEP. Unfolding was observed as an increase in fluorescence which was detected at an excitation wavelength of 492 nM and an emission wavelength of 610 nM. Data were plotted using GraphPad Prism and represent the average of at least three different measurements.

## Acknowledgement

This work utilized the cryo-electron Microscopy beamline CM01 at the European Synchrotron Radiation Facility, and the cryo EM-facility of the Julius-Maximilians-Universität Würzburg funded by the German Research Foundation (359471283, 456578072, 525040890). This research was supported by the German Research Foundation (KI 562 11-1) and the German Cancer Aid (70114277) to C.K.

## Data availability

Cryo EM data and coordinates have been deposited with EMDB and pdb, respectively. Class 1 and the XPD/DNA complex model are available under the accession codes EM-18206 and 8q7a. Class 2 data are available under EM-18206. Other research data will be made available upon request.

## Ethics declarations

The authors declare no competing interests.

## Author contributions

T.H., M.K., E.G., and S.M. carried out the biochemical studies. E.G. purified all proteins for this study. F.S. and E.G. generated XPD variants. H.N. produced the crosslinked DNA substrate. H.N. and C.H. were supervising crosslinked DNA substrate production. T.H. performed sample and cryo EM grid preparation. J.K. carried out data analysis and cryo EM data processing. J.K. and C.K. conceived the research project. J.K. and C.K. supervised experimental design and data interpretation. J.K. and C.K. were leading the manuscript writing process. All authors were involved in writing the paper and adhere to the ‘inclusion and ethics’ regulation.

## References

1 Compe, E. & Egly, J. M. TFIIH: when transcription met DNA repair. Nat Rev Mol Cell Biol 13, 343–354, doi:10.1038/nrm3350 (2012).

2 Kuper, J. et al. In TFIIH, XPD helicase is exclusively devoted to DNA repair. PLoS Biol 12, e1001954, doi:10.1371/journal.pbio.1001954 (2014).

3 Hoeijmakers, J. H. Genome maintenance mechanisms for preventing cancer. Nature 411, 366–374, doi:10.1038/35077232 (2001).

4 Gillet, L. C. & Scharer, O. D. Molecular mechanisms of mammalian global genome nucleotide excision repair. Chem Rev 106, 253–276, doi:10.1021/cr040483f (2006).

5 Scharer, O. D. Chemistry and biology of DNA repair. Angew Chem Int Ed Engl 42, 2946–2974, doi:10.1002/anie.200200523 (2003).

6 Scharer, O. D. Nucleotide excision repair in eukaryotes. Cold Spring Harb Perspect Biol 5, a012609, doi:10.1101/cshperspect.a012609 (2013).

7 Tsutakawa, S. E. et al. Envisioning how the prototypic molecular machine TFIIH functions in transcription initiation and DNA repair. DNA Repair (Amst) 96, 102972, doi:10.1016/j.dnarep.2020.102972 (2020).

8 Kuper, J. & Kisker, C. At the core of nucleotide excision repair. Curr Opin Struct Biol 80, 102605, doi:10.1016/j.sbi.2023.102605 (2023).

9 Diderich, K., Alanazi, M. & Hoeijmakers, J. H. Premature aging and cancer in nucleotide excision repair-disorders. DNA Repair (Amst) 10, 772–780, doi:10.1016/j.dnarep.2011.04.025 (2011).

10 Kokic, G. et al. Structural basis of TFIIH activation for nucleotide excision repair. Nat Commun 10, 2885, doi:10.1038/s41467-019-10745-5 (2019).

11 Kokic, G., Wagner, F. R., Chernev, A., Urlaub, H. & Cramer, P. Structural basis of human transcription-DNA repair coupling. Nature 598, 368–372, doi:10.1038/s41586-021-03906-4 (2021).

12 Kim, J. et al. Lesion recognition by XPC, TFIIH and XPA in DNA excision repair. Nature 617, 170–175, doi:10.1038/s41586-023-05959-z (2023).

13 van Eeuwen, T. et al. Cryo-EM structure of TFIIH/Rad4-Rad23-Rad33 in damaged DNA opening in nucleotide excision repair. Nat Commun 12, 3338, doi:10.1038/s41467-021-23684-x (2021).

14 Kappenberger, J. et al. How to limit the speed of a motor: the intricate regulation of the XPB ATPase and translocase in TFIIH. Nucleic Acids Res 48, 12282–12296, doi:10.1093/nar/gkaa911 (2020).

15 Neitz, H., Bessi, I., Kuper, J., Kisker, C. & Hobartner, C. Programmable DNA Interstrand Crosslinking by Alkene-Alkyne [2 + 2] Photocycloaddition. J Am Chem Soc 145, 9428–9433, doi:10.1021/jacs.3c01611 (2023).

16 Greber, B. J., Toso, D. B., Fang, J. & Nogales, E. The complete structure of the human TFIIH core complex. Elife 8, doi:10.7554/eLife.44771 (2019).

17 Wood, R. D. Mammalian nucleotide excision repair proteins and interstrand crosslink repair. Environ Mol Mutagen 51, 520–526, doi:10.1002/em.20569 (2010).

18 Cheng, K. & Wigley, D. B. DNA translocation mechanism of an XPD family helicase. Elife 7, doi:10.7554/eLife.42400 (2018).

19 Peissert, S. et al. In TFIIH the Arch domain of XPD is mechanistically essential for transcription and DNA repair. Nat Commun accepted (2020).

20 Fishburn, J., Tomko, E., Galburt, E. & Hahn, S. Double-stranded DNA translocase activity of transcription factor TFIIH and the mechanism of RNA polymerase II open complex formation. Proc Natl Acad Sci U S A 112, 3961–3966, doi:10.1073/pnas.1417709112 (2015).

21 Aibara, S., Schilbach, S. & Cramer, P. Structures of mammalian RNA polymerase II pre-initiation complexes. Nature 594, 124–128, doi:10.1038/s41586-021-03554-8 (2021).

22 Schilbach, S. et al. Structures of transcription pre-initiation complex with TFIIH and Mediator. Nature 551, 204–209, doi:10.1038/nature24282 (2017).

23 Gregersen, L. H. & Svejstrup, J. Q. The Cellular Response to Transcription-Blocking DNA Damage. Trends Biochem Sci 43, 327–341, doi:10.1016/j.Mbs.2018.02.010 (2018).

24 Lindsey-Boltz, L. A. et al. Nucleotide excision repair in Human cell lines lacking both XPC and CSB proteins. Nucleic Acids Res 51, 6238–6245, doi:10.1093/nar/gkad334 (2023).

25 Li, C. L. et al. Tripartite DNA Lesion Recognition and Verification by XPC, TFIIH, and XPA in Nucleotide Excision Repair. Mol Cell 59, 1025–1034, doi:10.1016/j.molcel.2015.08.012 (2015).

26 Selby, C. P., Lindsey-Boltz, L. A., Li, W. & Sancar, A. Molecular Mechanisms of Transcription-Coupled Repair. Annu Rev Biochem 92, 115–144, doi:10.1146/annurev-biochem-041522-034232 (2023).

27 Mietus, M. et al. Crystal structure of the catalytic core of Rad2: insights into the mechanism of substrate binding. Nucleic Acids Res 42, 10762–10775, doi:10.1093/nar/gku729 (2014).

28 Mathieu, N., Kaczmarek, N., Ruthemann, P., Luch, A. & Naegeli, H. DNA quality control by a lesion sensor pocket of the xeroderma pigmentosum group D helicase subunit of TFIIH. Curr Biol 23, 204–212, doi:10.1016/j.cub.2012.12.032 (2013).

29 Mathieu, N., Kaczmarek, N. & Naegeli, H. Strand- and site-specific DNA lesion demarcation by the xeroderma pigmentosum group D helicase. Proc Natl Acad Sci U S A 107, 17545–17550, doi:10.1073/pnas.1004339107 (2010).

30 Li, M. Z. & Elledge, S. J. Harnessing homologous recombination in vitro to generate recombinant DNA via SLIC. Nat Methods 4, 251–256, doi:10.1038/nmeth1010 (2007).

31 Wang, W. & Malcolm, B. A. Two-stage PCR protocol allowing introduction of multiple mutations, deletions and insertions using QuikChange Site-Directed Mutagenesis. Biotechniques 26, 680–682 (1999).

32 Kandiah, E. et al. CM01: a facility for cryo-electron microscopy at the European Synchrotron. Acta Crystallogr D Struct Biol 75, 528–535, doi:10.1107/S2059798319006880 (2019).

33 Zheng, S. Q. et al. MotionCor2: anisotropic correction of beam-induced motion for improved cryo-electron microscopy. Nat Methods 14, 331–332, doi:10.1038/nmeth.4193 (2017).

34 Punjani, A., Rubinstein, J. L., Fleet, D. J. & Brubaker, M. A. cryoSPARC: algorithms for rapid unsupervised cryo-EM structure determination. Nat Methods 14, 290–296, doi:10.1038/nmeth.4169 (2017).

35 Adams, P. D. et al. PHENIX: a comprehensive Python-based system for macromolecular structure solution. Acta Crystallogr D Biol Crystallogr 66, 213–221, doi:10.1107/S0907444909052925 (2010).

36 Burnley, T., Palmer, C. M. & Winn, M. Recent developments in the CCP-EM soqware suite. Acta Crystallogr D Struct Biol 73, 469–477, doi:10.1107/S2059798317007859 (2017).

37 Wood, C. et al. Collaborative computational project for electron cryo-microscopy. Acta Crystallogr D Biol Crystallogr 71, 123–126, doi:10.1107/S1399004714018070 (2015).

